# The Short Isoform of the Mouse Actin Adaptor Protein Synaptopodin-2 Activates Actin-Responsive Transcription Factors and Enhances Myoblast Fusion

**DOI:** 10.1101/2025.03.21.644618

**Authors:** Nandini Nagarajan Margam, FuiBoon Kai, Victor Chichkov, Colin Crist, Roy Duncan

**Affiliations:** Department of Microbiology and Immunology, Dalhousie University, Halifax, Nova Scotia, Canada, B3H4R2; Department of Human Genetics, McGill University, Montreal, Quebec, Canada, H3T1E2; Department of Biochemistry and Molecular Biology, Dalhousie University, Halifax, Nova Scotia, Canada, B3H4R2; Department of Pediatrics, Dalhousie University, Halifax, Nova Scotia, Canada, B3H4R2

## Abstract

Expression of synaptopodin-2 (SYNPO2), an actin cytoskeleton adaptor protein, is up-regulated following myoblast differentiation into myocytes and subsequently myotubes but no functional role in muscle development has been attributed to SYNPO2. We now show that of the three known mouse SYNPO2 isoforms, only the shortest isoform, SYNPO2As, is upregulated following differentiation of C2C12 myoblasts and primary mouse muscle satellite cells. Consistent with the differentiation-dependent expression pattern of these different isoforms, ectopic expression of SYNPO2As significantly increased myotube formation while the two large SYNPO2 isoforms (SYNPO2A or SYNPO2B) inhibited myocyte fusion into multinucleated myotubes. Irrespective of the fusion phenotype, all three isoforms enhanced migration of differentiating myoblasts. Knockdown studies using shRNA confirmed a pro-myogenic role for the short SYNPO2As isoform. Interestingly, SYNPO2As-transduced cells increased the transcript levels of late myogenic differentiation markers in the STARS (striated muscle activator of Rho signaling) and SRF (serum response factor) pathways that correlated with the enhanced fusion phenotype. These results identify the actin adaptor protein SYNPO2As as a new pro-myogenic factor that upregulates late myogenic differentiation markers and myotube formation.

## Introduction

Myogenesis is a tightly regulated, multi-step process whereby undifferentiated progenitor cells commit to form mononucleated myoblasts which then differentiate into elongated myocytes. Myocytes migrate and then align, adhere and fuse to one another to form nascent myotubes with several nuclei, which then fuse to each other and with additional myocytes to generate large, multinucleated mature myotubes (Abmayr and Pavlath 2012). Skeletal muscle regeneration also involves cell-cell fusion, where activated satellite cells (adult muscle stem cells) generate myogenic progenitors that fuse with damaged myotubes to regenerate myofibrils (Tierney and Sacco 2016). Studies in humans and multiple animal models have identified numerous pathways and proteins that control myogenesis. These pro-myogenic proteins include numerous transcription factors that drive the differentiation program (Singh and Dilworth 2013), and two recently identified muscle-specific membrane proteins required for myocyte fusion during muscle development and regeneration (Millay, O’Rourke et al. 2013, Quinn, Goh et al. 2017). Myogenesis is also highly reliant on extensive remodelling of the actin cytoskeleton to regulate cell migration, alignment and fusion (Kim and Chen 2019). While multiple proteins involved in cell-adhesion and actin remodelling have been identified as key players in myotube formation (Kim, Jin et al. 2015, Schejter 2016), it seems likely that additional actin regulators important for myogenesis remain to be identified.

Synaptopodin-2 (SYNPO2) is an actin regulator and mechanosensor involved in maintaining muscle tissue homeostasis. SYNPO2, also known as myopodin or chicken fesselin, belongs to the podin protein family and shares 48% sequence identity to the founding member of the family, synaptopodin-1 (SYNPO1) (Weins, Schwarz et al. 2001, Chalovich and Schroeter 2010). SYNPO1 and SYNPO2 are both proline-rich actin binding proteins that modulate cell shape and motility. SYNPO2 co-localizes with α-actinin and filamin in a punctuated staining pattern along actin filaments in skeletal and smooth muscle (Weins, Schwarz et al. 2001, Linnemann, van der Ven et al. 2010), and SYNPO2 expression is linked to actin polymerization in smooth muscle cells (Turczynska, Sward et al. 2015). SYNPO2 has also been shown to promote *in vitro* actin nucleation, polymerization and bundling into linear actin fibers (Beall and Chalovich 2001, Schroeter and Chalovich 2005, Linnemann, Vakeel et al. 2012, Schroeter, Orlova et al. 2013). SYNPO2 also functions as a scaffold protein in smooth muscle cells to assemble a mechanosensor complex that activates a selective autophagy pathway in response to tension in the actin cytoskeleton, and the short isoform, SYNPO2e, is required for myofibril stabilization (Ulbricht, Eppler et al. 2013, Lohanadan, Molt et al. 2021). While SYNPO2 is clearly an actin regulator and mechanosensor involved in differentiated muscle cell homeostasis, it has no known role in myogenesis and myotube formation.

Most of our early understanding of how SYNPO2 affects actin dynamics in cells derives from studies in cancer cells where loss of SYNPO2 expression, due to gene deletion or methylation-dependent epigenetic silencing, strongly correlates with increased tumour cell invasion in prostate, bladder and colon cancer (Lin, Yu et al. 2001, De Ganck, De Corte et al. 2008, Esteban, Moya et al. 2012). Dysregulation of SYNPO2 variants has also been linked to the pathogenesis of nephrotic syndrome via Rac1-ARP3 dysregulation (Mao, Schneider et al. 2021). The human *SYNPO2* gene has seven exons that are differentially spliced to generate four isoforms (SYNPO2A-D) (De Ganck, De Corte et al. 2008, Kai and Duncan 2013). Human SYNPO2A-C share a common N-terminus but have unique C-termini, while SYNPO2D has the same C-terminus as SYNPO2C but a unique N-terminus (Fig. 1A). Transcription initiation from an internal promoter in exon 4 and translation initiation at an in-frame methionine codon near the beginning of exon 5 generates a fifth isoform, SYNPO2As (Linnemann, van der Ven et al. 2010), which is a shorter, N-terminally truncated version of SYNPO2A (Fig. 1A). The conserved exon 5 region present in all five SYNPO2 isoforms contains multiple actin binding sites, and binding sites for several proteins that regulate actin or focal adhesion dynamics and chaperone-assisted selective autophagy (Yu and Luo 2006, Linnemann, van der Ven et al. 2010, Yu and Luo 2011, Linnemann, Vakeel et al. 2012, Ulbricht, Eppler et al. 2013)). All five of these isoforms can promote or inhibit migration of prostate cancer cells, depending on the nature of the external migration stimulus. These motility responses reflect the effects of SYNPO2As on the Rho/ROCK signaling pathway, a central regulator of actin dynamics and cell motility. For example, in response to serum stimulation of prostate cancer PC3 cells, SYNPO2As activates RhoA and uses a ROCK-dependent pathway to promote assembly of parallel actin bundles and nascent focal adhesions at the leading cell edge, resulting in increased lamellipodia formation and enhanced non-directional cell migration (Kai and Duncan 2013, Kai, Fawcett et al. 2015). It is unknown whether SYNPO2 isoforms exert similar effects on the actin cytoskeleton and/or migration of muscle cells.

**Figure 1:**
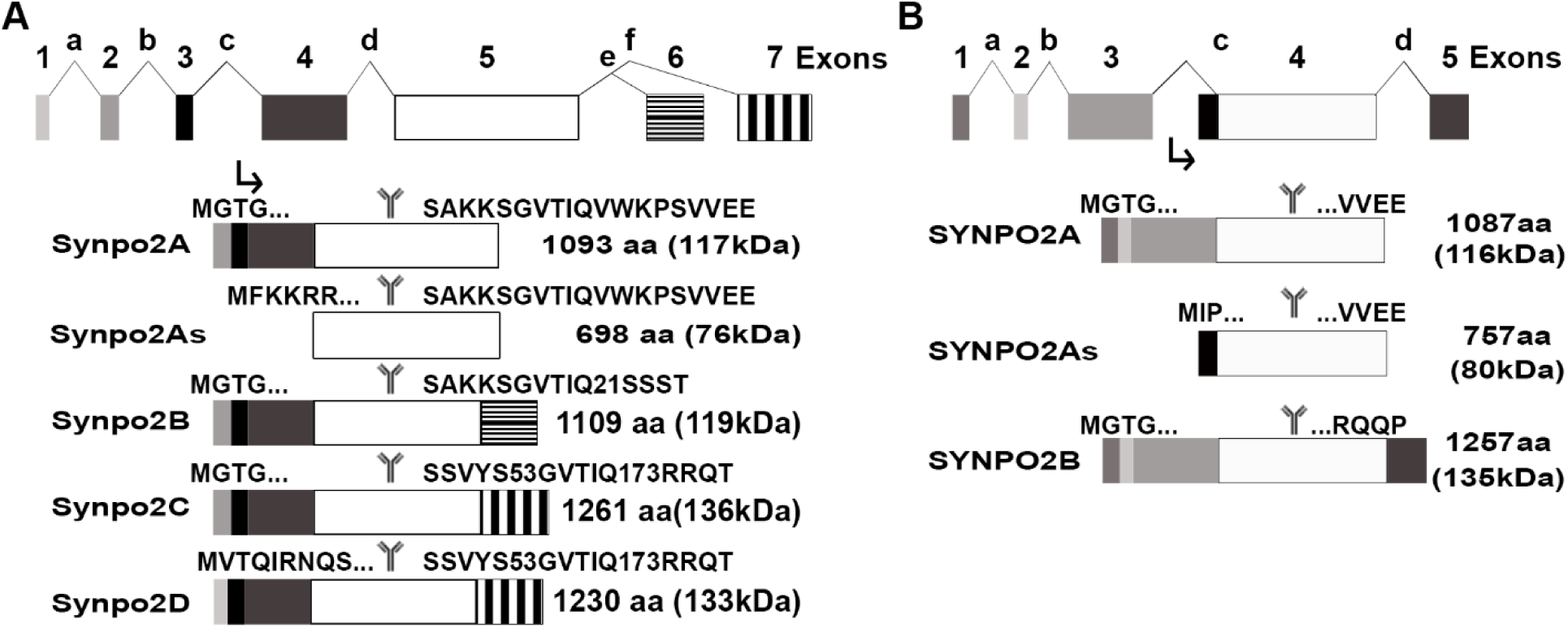
Human and mouse SYNPO2 isoforms. Exon arrangements of the human (A) and mouse (B) SYNPO2 genes and protein isoforms. The protein isoforms are translated from alternatively spliced mRNAs generated from an upstream promoter, or in the case of the short isoforms (As suffix) from an internal promoter (downward arrow in each panel). Upper panels depict the gene, with exons indicated by numbered shaded rectangles and introns by alphabetically labelled chevrons. Amino acid sequences above each protein isoform indicate the N- and C-terminal sequences. The number of amino acid residues and predicted molecular mass (in brackets) for each isoform are indicated. The centrally located conserved exon (white rectangle) contains the epitope recognized by the commercial antiserum (indicated by the Y symbol) and several protein interaction motifs described in the text.

Most studies on mouse SYNPO2 used the 757 amino acid isoform of SYNPO2, a homologue of human SYNPO2As (Fig. 1B). The transcript for this short mouse SYNPO2As isoform presumably derives from an internal promoter contained within exon 3 or intron c (Fig. 1B), like the internal promoter responsible for transcription of the human SYNPO2As isoform (Linnemann, van der Ven et al. 2010). The NCBI and Ensemble databases report additional mouse SYNPO2 isoforms derived by alternate splicing, homologues of the longer human SYNPO2A-C isoforms (Fig. 1A and B). Similar to the human isoforms, these longer mouse isoforms share a common N-terminus but possess unique C-termini, and the shorter SYNPO2As isoform has the same C-terminus as SYNPO2A but is truncated at the N-terminus (Fig. 1B). However, mouse SYNPO2As contains a unique 17-residue N-terminus derived by translation initiation from an in-frame methionine start codon contained within intron c. Aside from the involvement of the larger isoforms in chaperone-assisted selective autophagy (Ulbricht, Eppler et al. 2013), there are no other studies describing roles for the various murine SYNPO2 isoforms.

Human SYNPO2 expression is upregulated shortly after inducing myoblast differentiation (Linnemann, van der Ven et al. 2010) suggesting SYNPO2 likely serves an important, though undefined, role during myogenesis. To address this knowledge deficit, SYNPO2 expression was manipulated in differentiating C2C12 mouse myoblasts and the effects on myoblast migration, differentiation and myotube formation were examined. Results indicated the mouse SYNPO2As isoform, but not the larger isoforms involved in chaperone-assisted autophagy, functions as a pro-myogenic factor that activates the SRF and STARS pathways to upregulate muscle-specific transcripts and enhance myoblast fusion.

## RESULTS

### SYNPO2 isoforms are differentially expressed following C2C12 myoblast differentiation

To detect expression of endogenous SYNPO2 isoforms in myoblast cells pre- and post-differentiation, sub-confluent mouse C2C12 myoblasts were differentiated by culturing in medium containing 2% horse serum. As seen in Giemsa-stained microscopic images of cells (Fig. 2A), mononucleated myoblasts elongated into spindle-shaped cells and started to align next to each other by 1-2 days post-induction of differentiation (dpi), and numerous cells fused to form multinucleated myotubes by 3-4 dpi. Western blotting using a commercial polyclonal antibody raised against the conserved exon present in all mouse SYNPO2 isoforms revealed a progressive increase in the appearance of an 80 kDa polypeptide starting at 1 dpi (Fig. 2B), which corresponded to the predicted molecular weight of the 757-residue SYNPO2As isoform (Fig. 1B). There was no detectable differentiation-dependent upregulated expression of polypeptides migrating in the expected range for the larger SYNPO2 isoforms (Fig. 2B), whose predicted molecular weights range from 116-135 kDa (Fig. 1B).

**Figure 2:**
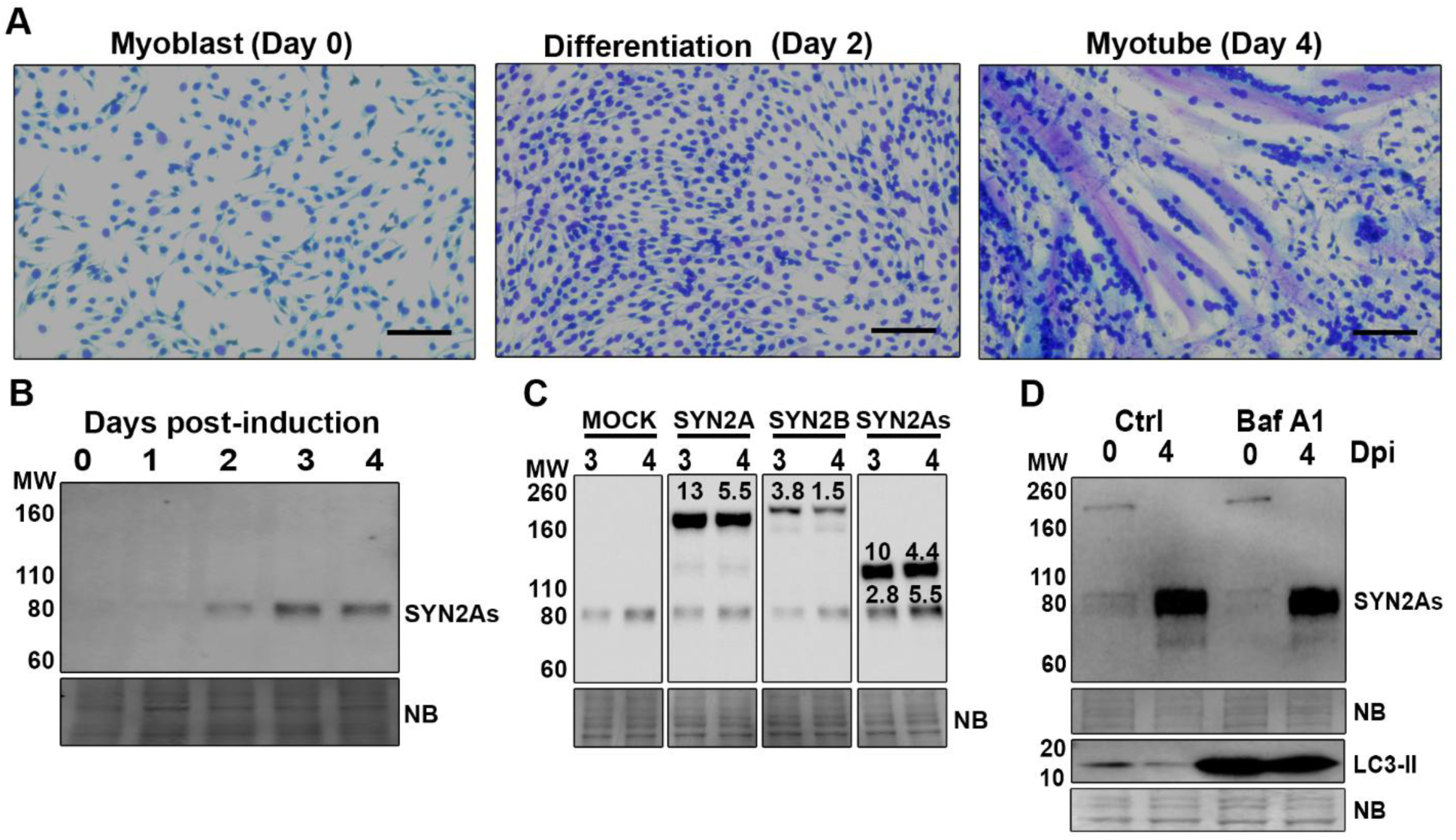
SYNPO2As expression is upregulated following myoblast differentiation. (A) Representative microscopic images of Giemsa-stained C2C12 mouse myoblasts cultured in growth medium (Day 0) or following 2 or 4 days growth in differentiation medium. Scale bars = 2µm. (B) Western blot of C2C12 cell lysates collected at 1-4 dpi and probed with anti-SYNPO2. (C) Western blot of C2C12 cell lysates from transduced cells stably expressing the three mouse SYNPO2 isoforms (SYN2A, SYN2B and SYN2As) collected at 3 or 4 dpi and probed with anti-SYNPO2. (D) C2C12 cells at 0 or 4 dpi were treated with DMSO (Ctrl) or bafilomycin A1 (BafA1) to inhibit autophagy, and western blots of cell lysates were probed with antibodies specific for SYNPO2 (SYN2As) or LC3-II as a marker of autophagic flux. Molecular weight markers in kDa (MW) are indicated on the left of each blot and naphthol blue (NB) stained membranes were used as loading controls.

To determine whether the anti-SYNPO2 antibody could recognize the larger SYNPO2 isoforms, cDNAs were obtained using mRNAs isolated from proliferating, undifferentiated mouse C2C12 myoblast cells and terminal primers specific for the two long isoforms, SYNPO2A and SYNPO2B. Western blots of cell lysates from stably transduced C2C12 cells constitutively expressing cDNAs of the long and short SYNPO2 isoforms revealed all three isoforms were easily detected using the anti-SYNPO2 antiserum (Fig. 2C). Interestingly, ectopically expressed SYNPO2As displayed an unusual gel migration pattern, generating the predicted 80 kDa polypeptide observed for endogenous SYNPO2As and a predominant 110 kDa polypeptide (Fig. 2C), the ratios of which varied considerably between experiments (data not shown). Slower migrating species of SYNPO2As have been previously reported but unexplained (Weins, Schwarz et al. 2001, Schroeter, Beall et al. 2008, Turczynska, Sward et al. 2015). The fold change in expression levels of the ectopically expressed SYNPO2As isoform (combined 110 kDa and 80 kDa polypeptides) relative to the endogenous 80 kDa polypeptide was approximately 10-12 fold. (Fig. 2C). The longer SYNPO2 isoforms also displayed aberrant migration, migrating in the vicinity of the 160 kDa marker, considerably slower than expected based on their predicted molecular weights. Human SYNPO2 isoforms in cancer cells were previously noted to migrate aberrantly slow (De Ganck, De Corte et al. 2008). The retarded gel mobility of SYNPO2 isoforms may reflect their proline-rich nature that suggests SYNPO2 is an intrinsically disordered protein, which are known to migrate slower on SDS-PAGE gels (Iakoucheva, Kimzey et al. 2001).

The ability to obtain cDNAs from undifferentiated C2C12 cells indicates the mRNAs for all three SYNPO2 isoforms were present in these cells. On overexposed western blots (Fig. 2D), an 80 kDa band comigrating with SYNPO2As was faintly detectible in undifferentiated cells. The larger SYNPO2 isoforms are reported to be degraded by selective autophagy in differentiated C2C12 myoblasts (Ulbricht et al., 2013). Treatment of C2C12 cells pre- and post-differentiation with bafilomycin A1 inhibited lysosome acidification and autophagic degradation, as evident by increased detection of LC3-II (a marker of autophagic flux) but did not result in detection of the larger SYNPO2 isoforms (Fig. 2D). Similar results were reported in human skeletal myoblasts, where no SYNPO2 isoforms were detectable in undifferentiated myoblasts and only SYNPO2As displayed differentiation-dependent expression (Linnemann, van der Ven et al. 2010). Thus, while SYNPO2 isoforms may be expressed at very low levels in undifferentiated mouse myoblasts, only SYNPO2As is significantly upregulated shortly after inducing myoblast differentiation.

### Differentiation-dependent expression of SYNPO2As in mouse satellite cells

Satellite cells obtained from adult mouse muscle were used to confirm the expression pattern of SYNPO2 isoforms in primary cells. Satellite cells are adult muscle stem cells required for post-natal regeneration of muscle. Normally mitotically quiescent, satellite cells activate cell cycle and the myogenic program in response to injury and are the source of additional myonuclei for damaged muscle fibers following cell-cell fusion (Tierney and Sacco 2016). Satellite cells from the diaphragm and abdominal muscles of adult *Pax3^GFP/+^* mice were isolated by flow cytometry, where both paired box protein Pax-3 (PAX3) and paired box protein Pax-7 (PAX7) paired-homeodomain transcription factors commonly mark satellite cells (Montarras, Morgan et al. 2005). In immunofluorescence studies, we combined antibodies against SYNPO2 with Pax7, myoblast determination protein (MYOD) and troponin T to detect SYNPO2 expression as fresh isolated satellite cells (Day 0, Pax7 positive) enter the myogenic program (Day 3, MyoD positive) and differentiate (Day 5, troponin T positive) during culture. At day 0, SYNPO2 expression was barely detectable in satellite cells by immunofluorescence or by Western blotting (Fig. 3A and B). Increased detection of SYNPO2 in the cytoplasm was apparent after 3 days in culture, a timepoint when the transcription factor MyoD was detectable in most cells (Fig. 3A). By 5 days in culture, Troponin T- and SYNPO2-positive multinucleated myotubes were apparent (Fig. 3A), with SYNPO2 localizing predominantly in the cytoplasm in a parallel, linear staining pattern, as previously reported along actin filaments in primary human skeletal muscle cells (Linnemann, van der Ven et al. 2010). Western blotting revealed a similar pattern of upregulated SYNPO2 expression, with an 80 kDa polypeptide corresponding to SYNPO2As easily detectable following 5 days in culture, coincident with detection of MHC and MyoD (Fig. 3B). No evidence of upregulated SYNPO2A or SYNPOB expression was detectable by western blotting in the differentiated satellite cells. Hence, in both primary and continuous mouse cells the only SYNPO2 isoform whose expression detectibly increases following myoblast differentiation is the short SYNPO2As isoform.

**Figure 3:**
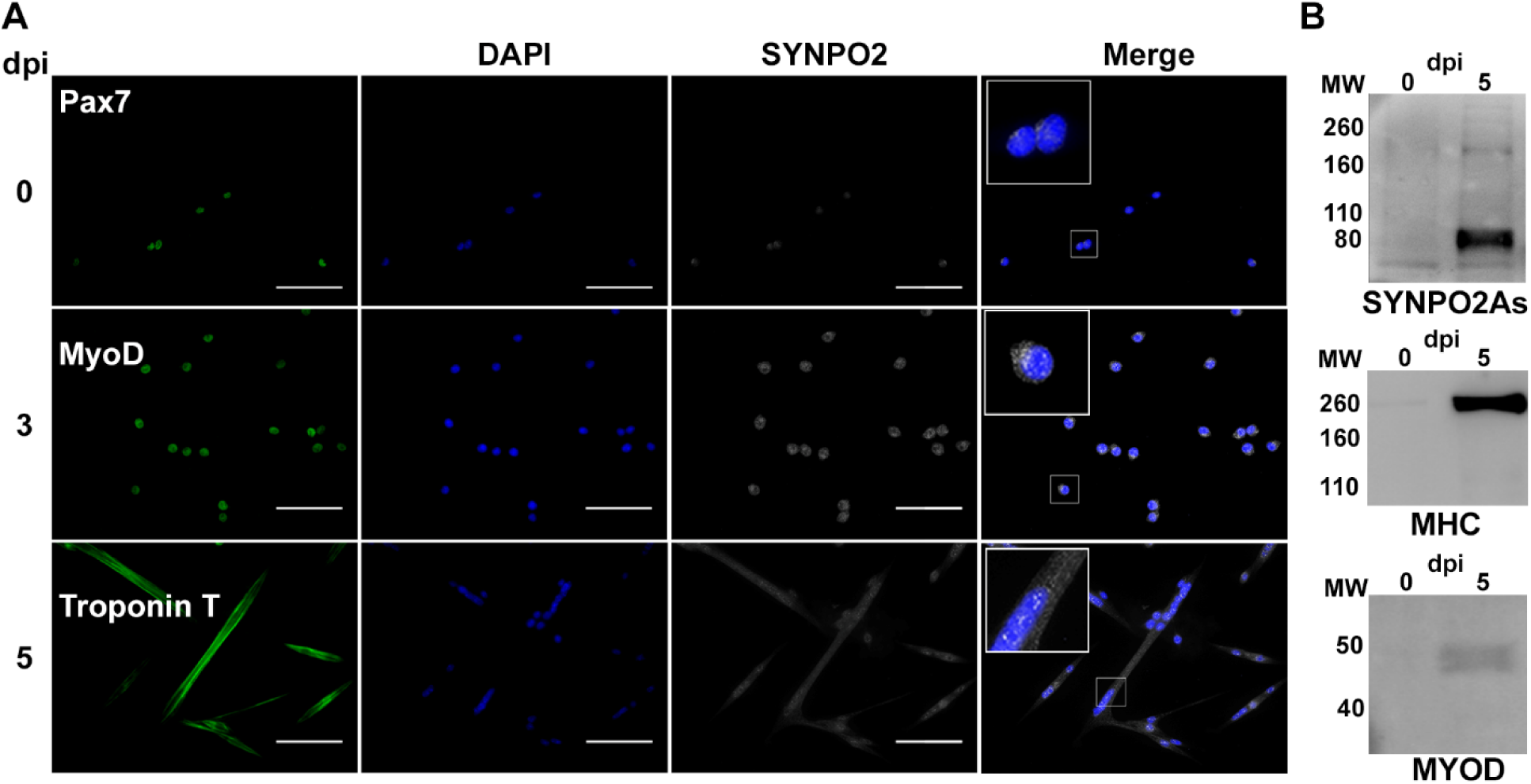
SYNPO2As expression is upregulated following differentiation of primary satellite cells. (A) Satellite cells from *Pax3^GFP/+^* mice were sorted and immunostained at 0, 3 or 5 dpi. Cells were immunostained with antibodies against SYNPO2 (black and white) and either Pax7, a satellite cell marker (0 dpi), MyoD, a myoblast differentiation marker (3 dpi), or Troponin T, a late myogenic marker (5 dpi) (all green). Nuclei were stained with DAPI (blue). Merge is an overlay of the DAPI and SYNPO2 images. (B) Western blots of *Pax3^GFP/+^* satellite cell lysates at 0 or 5 dpi and probed with antibodies against SYNPO2 (SYN2As), myosin heavy chain (MHC) or MyoD. Molecular weight markers in kDa (MW) are indicated on the left of each blot. Scale bars = 2µm.

### SYNPO2 isoforms differentially affect myotube formation

To determine the effects of SYNPO2 isoforms on myogenesis, C2C12 cells stably transduced with an empty retrovirus vector or with retrovirus vectors expressing the individual SYNPO2 isoforms were differentiated, and microscopic examination of Giemsa-stained monolayers was used to quantify the percent of nuclei in myotubes (Fig. 4A). Ectopic expression of SYNPO2A and SYNPO2B significantly delayed myotube formation at 4 dpi by ∼50% compared to mock transduced cells (Fig. 4B). Conversely, ectopic expression of SYNPO2As reproducibly increased myotube formation by ∼50% relative to mock-transduced cells. Immunofluorescence microscopy using the polyclonal anti-SYNPO2 antibody to detect endogenous SYNPO2As and anti-myc antibody to detect the ectopically expressed isoforms detected endogenous SYNPO2As in a punctuated staining pattern along linear actin filaments in cells co-stained with phalloidin to image actin filaments (Fig. 5), as previously shown in human skeletal muscle cells (Linnemann, van der Ven et al. 2010). Ectopically expressed SYNPO2As showed the same staining pattern as endogenous SYNPO2As, although more intense, consistent with its increased expression level. The larger SYNPO2A and SYNPO2B isoforms also localized along actin fibers (Fig. 5). Qualitatively, no gross change was observed in the actin cytoskeleton of differentiating myocytes expressing the various SYNPO2 isoforms (Fig. 5).

**Figure 4:**
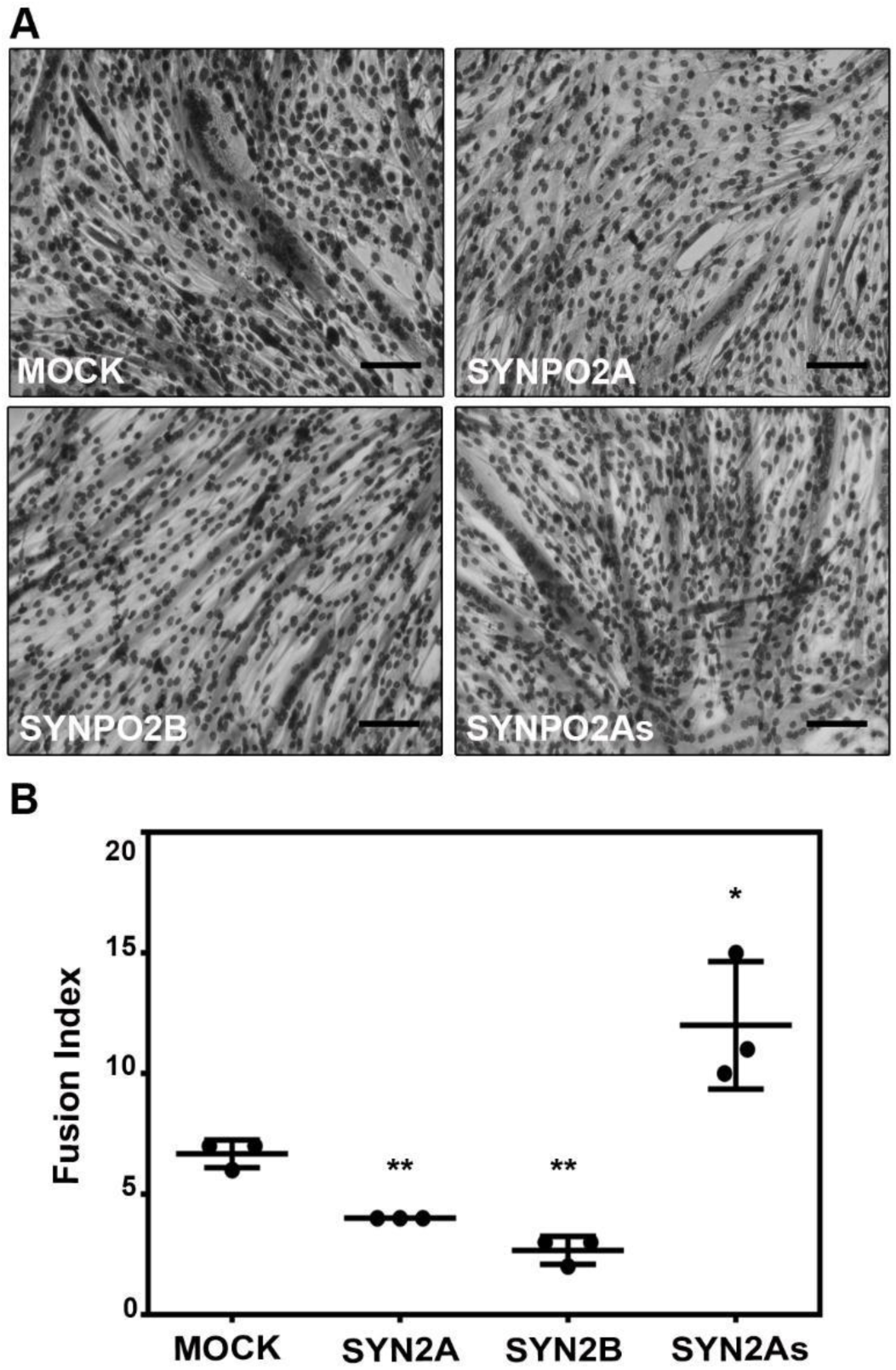
Differential effects of SYNPO2 isoforms on C2C12 myotube formation. (A) Giemsa-stained microscopic images of C2C12 cells stably expressing the indicated SYNPO2 isoforms at 4 dpi. Scale bars = 2µm. (B) C2C12 cells stably transduced with an empty retrovirus vector (Mock) or with retrovirus vectors expressing the indicated SYNPO2 isoforms were induced to differentiate, and the fusion index of cells at 4 dpi was quantified from the Giemsa-stained microscopic images. Results are presented as the mean ± SD of the fusion index (percent of nuclei present in syncytia) from triplicate samples in three independent experiments and statistical significance was determined using unpaired Student *t* test. Statistical significance: * p < 0.05, ** p < 0.01.

**Figure 5:**
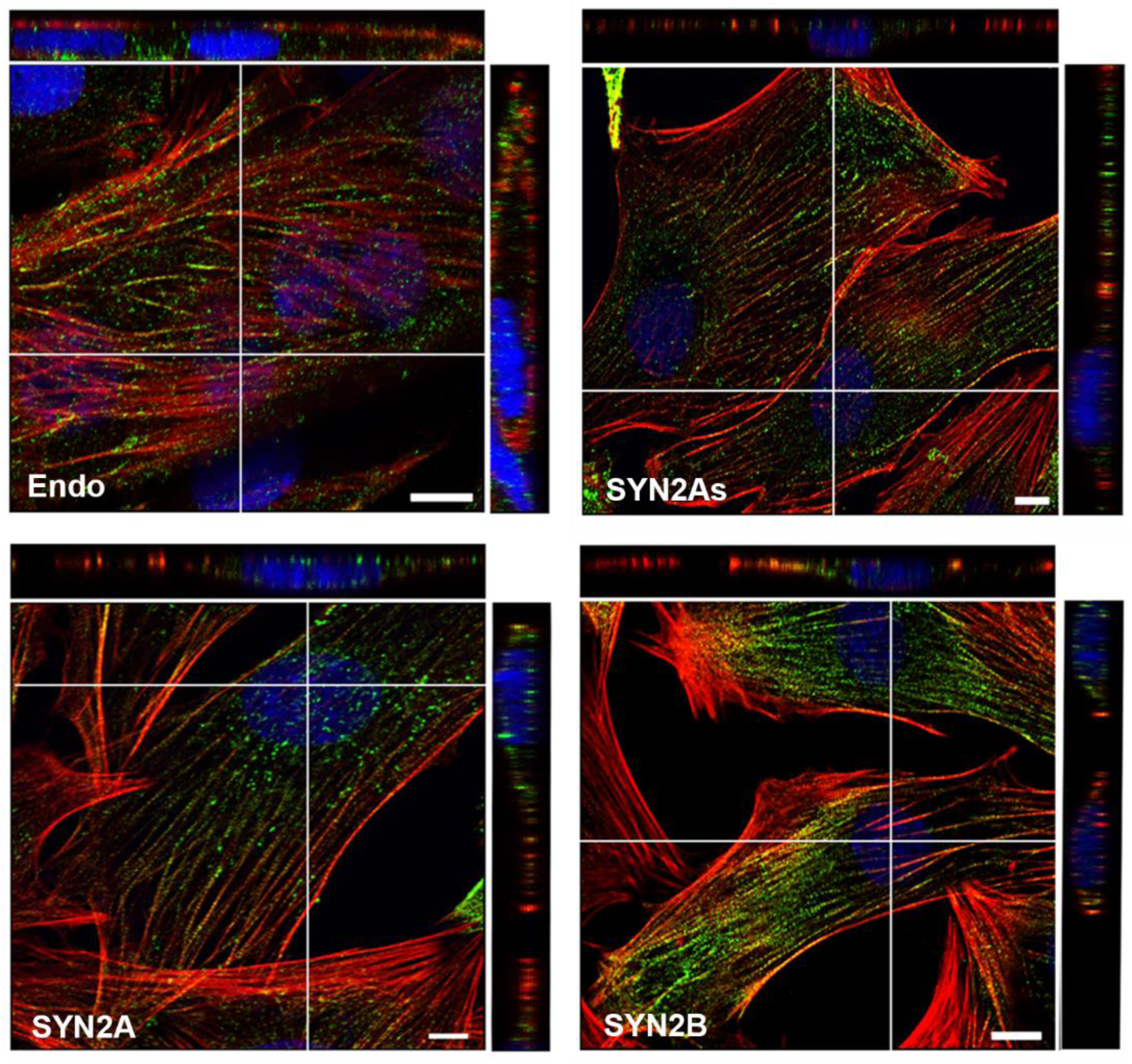
SYNPO2 isoforms associate with cytoplasmic actin filaments post-differentiation. Normal C2C12 cells expressing endogenous SYNPO2As (Endo) or C2C12 cells stably transduced with N-terminally myc-tagged SYNPO2 isoforms (SYN2As, SYN2A, SYN2B) were induced to differentiate for 2 dpi. Endogenous SYNPO2As was stained with anti-SYNPO2 antibody and cells ectopically expressing myc-tagged SYNPO2 isoforms were stained with anti-myc antibody and Alexa fluor 488-conjgated secondary antibodies (green). Filamentous actin was stained with Alexa-555 conjugated phalloidin (red) and nuclei were stained with DRAQ5 (blue). Images are one slice from a z-stack and orthogonal views (white lines depict the xz and yz slice) are shown on the top and right of each image. Scale bar = 10µm.

The ectopic SYNPO2 expression results were confirmed using shRNA-mediated knockdown of endogenous SYNPO2. Two shRNAs were designed, one targeting the unique N-terminus of SYNPO2As (shRNA1) and the second (shRNA2) targeting the conserved exon present in all the three isoforms (Fig. 1B). Since endogenous SYNPO2A and SYNPO2B were undetectable in C2C12 myoblasts (Fig. 1B), transiently transfected HEK293 cells expressing these two isoforms were used to assess the knockdown efficiency of shRNA2. As shown by western blotting (Fig 6A), shRNA2 almost completely inhibited expression of both SYNPO2A and SYNPOB. The knockdown efficiency of shRNA1 and shRNA2 on endogenous SYNPO2As was determined in differentiated C2C12 myoblasts at 3 dpi, which inhibited expression of SYNPO2As by ∼50% and ∼80% respectively (Fig. 6B). Inhibiting expression of endogenous SYNPO2As using shRNA1 significantly decreased the rate of myotube formation by ∼50% relative to cells expressing a non-targeting shRNA. This level of inhibition was not increased using shRNA2 to inhibit expression of all SYNPO2 isoforms (Fig. 6C), consistent with the lack of detectable differentiation-dependent expression of SYNPO2A and SYNPO2B (Fig. 2B). This is the first indication that SYNPO2As, but not SYNPO2A or SYNPO2B, functions as a pro-myogenic factor to promote myotube formation following myoblast differentiation.

**Figure 6:**
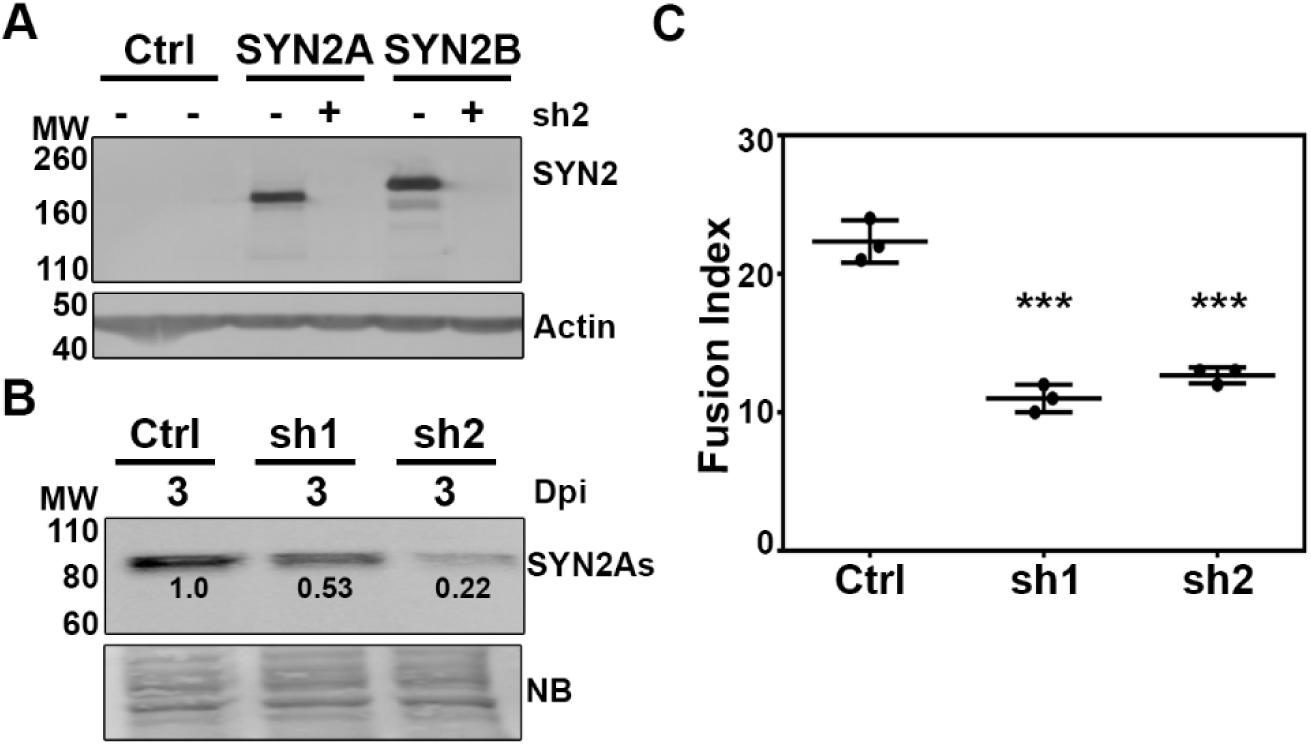
Knockdown of SYNPO2As inhibits myotube formation. (A) Western blot of cell lysates from HEK293 cells transiently transfected with empty plasmid (Ctrl) or with plasmids expressing SYNPO2A or SYNPO2B (SYN2A and SYN2B) and co-transfected with plasmids expressing the non-targeting shRNA (-) or expressing shRNA2 (sh2) that targets all three SYNPO2 isoforms, probed with anti-SYNPO2 (top panel) or actin. (B) Western blot of cell lysates from C2C12 cells at 3 dpi stably expressing shRNA1 or 2 (sh1, sh2) that target the regions encoding the unique N-terminus of SYNPO2As isoform or the conserved region present in all isoforms, respectively, or a non-targeting control shRNA (Ctrl), probed with anti-SYNPO2. Numbers indicate the fold change of the SYNPO2As polypeptides relative to cells transduced with the non-targeting shRNA at 3 dpi. Naphthol blue (NB) stained membrane was used as a loading control. Molecular weight markers (MW) in kDa are indicated on the left of each blot. (C) C2C12 cells stably expressing the shRNAs described in panel B were induced to differentiate and the fusion index of cells at 3 dpi was quantified from the Giemsa-stained microscopic images. Results are the mean fusion index ± SD from duplicate wells in three independent experiments and statistical significance was determined using unpaired Student *t* test. Statistical significance: *** p ≤ 0.005.

### SYNPO2As increases detection of late, but not early, markers of the myogenesis program

To determine whether SYNPO2As enhanced myoblast fusion by exerting an effect on the myogenic differentiation program, differentiation-dependent expression of several key factors in the myogenic program was examined by Western blotting. MYOD is a bHLH family transcription factor required to turn on the myoblast differentiation program (Tapscott 2005), and upregulates expression of a second essential bHLH family transcription factor, myogenin (MYOG) (Andres and Walsh 1996). MHC is an ATP-dependent motor protein involved in contracting actin filaments and is a marker of differentiated myocytes (Wells, Edwards et al. 1996). Antibodies specific for these differentiation markers were used to probe Western blots of transduced C2C12 cell lysates expressing SYNPO2As harvested at various days post-differentiation. As shown (Fig. 7A), there was no discernible qualitative difference in the kinetics or extent of upregulated expression of the early myogenic markers MYOD and MYOG in cells ectopically expressing SYNPO2As compared to mock-transduced cells, which was confirmed by quantification of the blots (Fig. 7D and E). In contrast, ectopic expression of SYNPO2As on 2 dpi and 3 dpi enhanced expression of the late differentiation MHC marker by 2-4-fold (Fig. 7C). Enhanced expression of MHC was not observed in cells expressing shRNA2 that inhibited endogenous SYNPO2 expression (Fig. 7B and Supplementary Figure 1). Thus, SYNPO2As increases the expression of late differentiation markers such as MHC, independent of any perturbation of the early and intermediate stages of the myogenesis differentiation program.

**Figure 7:**
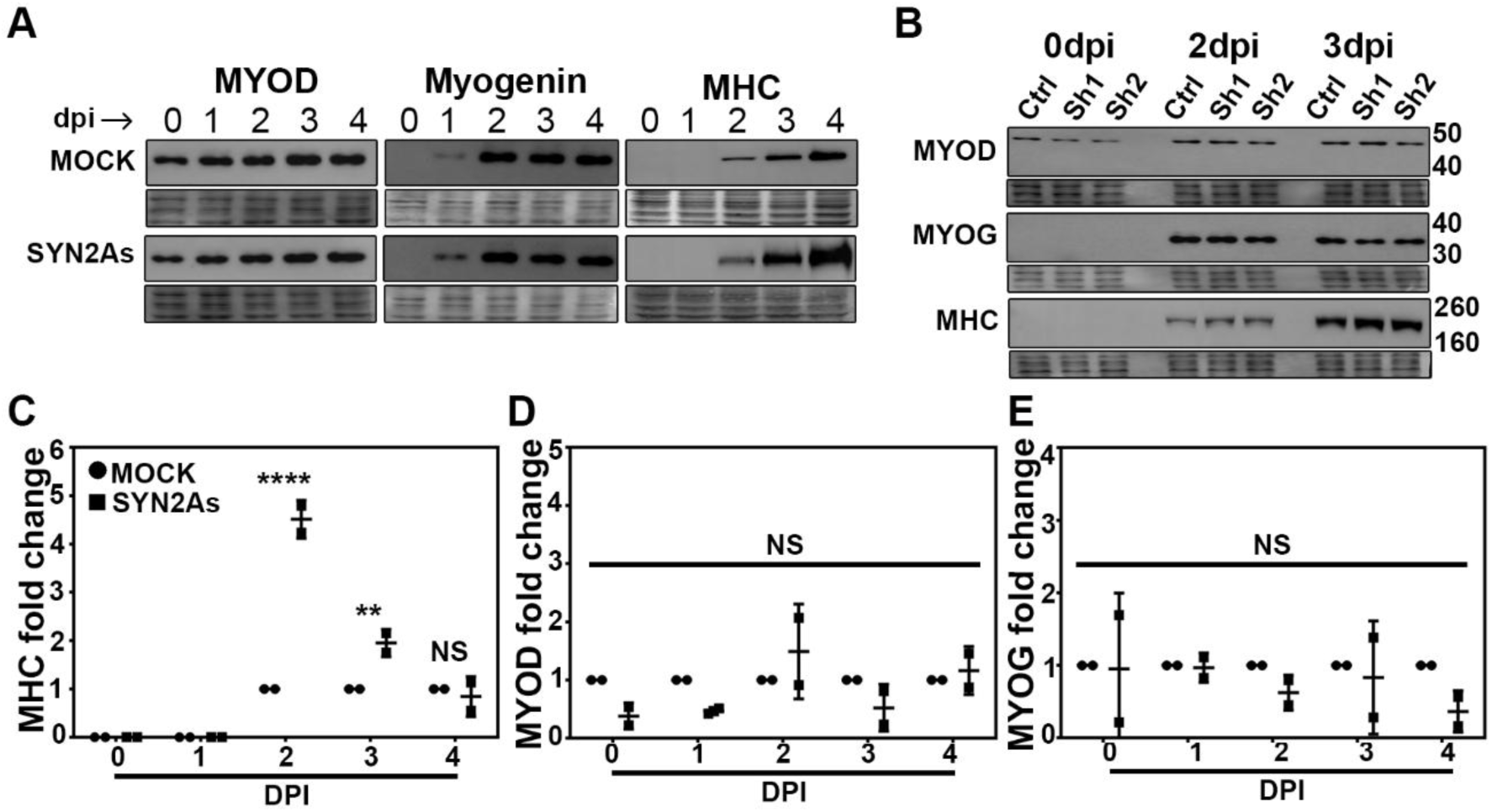
Knockdown or ectopic expression of SYNPO2 isoforms does not affect early differentiation. Western blots of C2C12 cells mock-transduced or transduced with retrovirus vector expressing SYNPO2As (SYN2As) (A) or stably expressing shRNA1 (Sh1) that targets SYNPO2As or shRNA2 (Sh2) that targets all SYNPO2 isoforms (B). Lysates were harvested at the indicated dpi and blots were probed with antibodies specific for MyoD, myogenin or myosin heavy chain (MHC). Molecular weight markers in kDa are indicated on the right, and naphthol blue stained blots (lower panels below each of the antibody probed blots) were used as a loading control. (C-E) Quantified data of MHC (C), MYOD (D) and MYOG (E) Western blot represented as the mean ± SD from two independent experiments of mock transduced and SYNPO2As expressing cell lysates collected at 0-4 dpi. Statistical significance was determined using two-way ANOVA with post-hoc by Bonferroni. Statistical significance: **p value < 0.01, ****p value < 0.001, and NS = Not significant.

### SYNPO2As-transduced cells increase myogenic markers with contractile function

In smooth muscle cells, SYNPO2 is a downstream target of serum response factor (SRF), and knocking down SYNPO2 in these cells decreases the transcript levels of late smooth muscle differentiation markers such as dystrophin and SM22α (Turczynska, Sward et al. 2015). The SRF pathway is regulated by the ratio of soluble:filamentous actin and it has been suggested that SYNPO2 utilizes its actin polymerization property to regulate this pathway (Turczynska, Sward et al. 2015). Since we observed a significant increase in MHC protein level when SYNPO2As was overexpressed in cells and these cells enhanced myoblast fusion, we wanted to determine whether SYNPO2As also triggered the expression of late differentiation markers in skeletal muscle cells. We therefore performed RT-qPCR on RNA extracted from control- or SYNPO2As-transduced C2C12 cells on days 0 and 3 post-differentiation using probes specific for selected genes from the SRF, striated muscle activator of Rho signalling (STARS), and forkhead box protein O1 (FOXO1), also known as forkhead in rhabdomyosarcoma (FKHR), pathways. These three pathways are well-known for regulating muscle-specific transcripts (Bois and Grosveld 2003, Miano, Long et al. 2007, Wallace, Della Gatta et al. 2016, Randrianarison-Huetz, Papaefthymiou et al. 2018). SYNPO2As significantly enhanced transcript levels of *MyoG*, *Srf*, *Stars*, α-actin (*Acta1*), mitochondrial 2 (*Ckmt2*), slow-type myosin heavy chain (*MyHC-I*) and all fast-type myosin heavy chains (*MyHC-IIa*, *IIb* and *IIx*), β-myosin heavy chain (*Myh7*) and tenascin-C (*Tnc*), but not *MyoD*, myogenic factor 5 (*Myf5*), prosaposin (*Pasp*) and frizzled-4 (*Fdz4*) (Fig. 8). These results were largely confirmed by quantifying the mRNA levels of the above-mentioned transcripts in SYNPO2 knockdown cells by RT-qPCR. Knocking down SYNPO2 significantly reduced the transcript level of genes that were upregulated in the SYNPO2As-transduced cells, such as *Acta1, MyHCIIb,* and *MyHCIIx* (Supplementary Fig. 2). The transcript levels of other genes that were upregulated in the SYNPO2As-transduced cells, such as *Srf*, *Stars* and *MyoG* (Supplementary Fig. 2) were not reduced significantly, but there was a downward trend in the mean expression level of these markers one day post-differentiation. These preliminary results need to be replicated and extended to protein expression levels to be conclusive, but the data are consistent with the hypothesis that SYNPO2As utilizes its actin polymerizing activity to regulate transcription of differentiation markers tbat promote early myotube formation.

**Figure 8:**
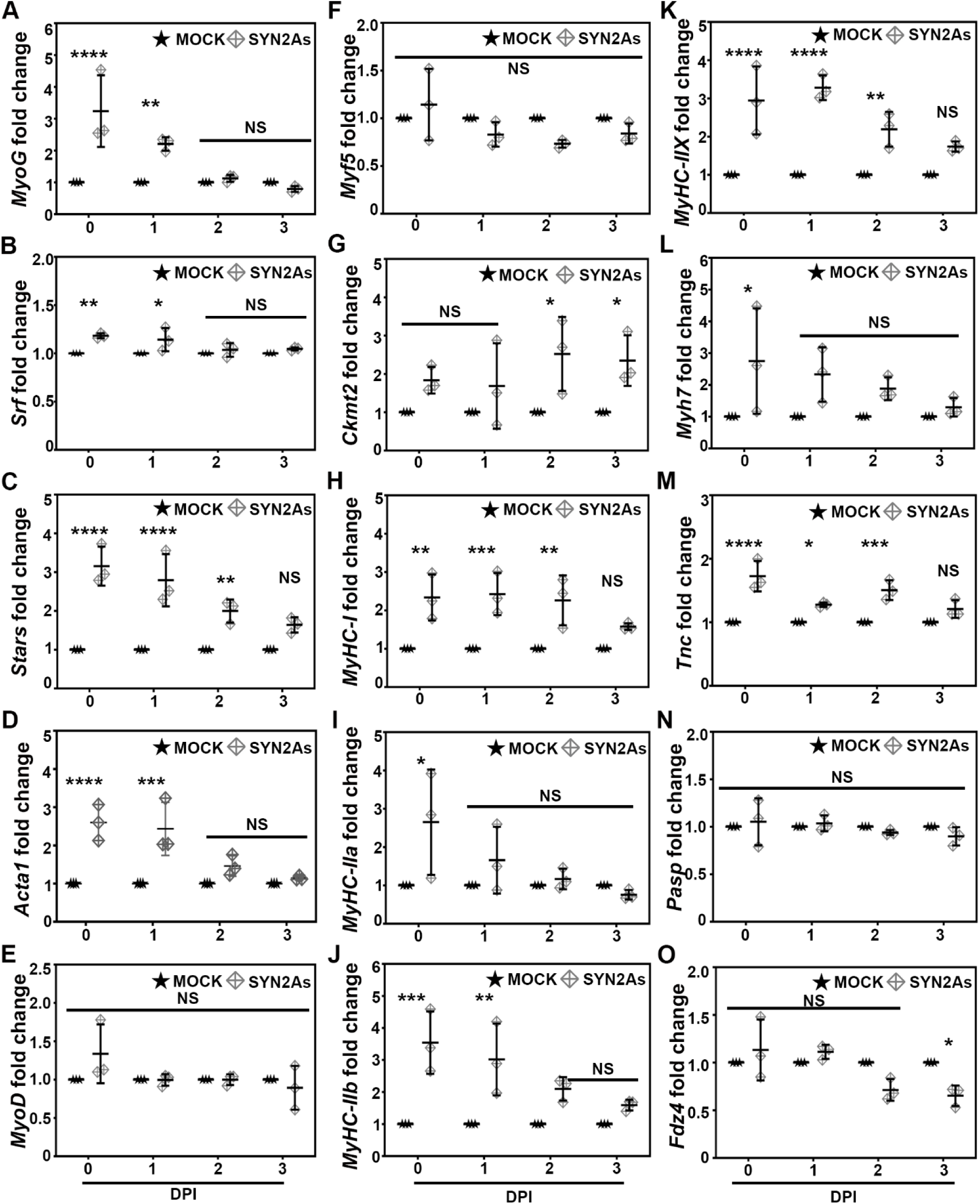
SYNPO2As overexpression increased selective muscle specific transcripts. Quantified data of mRNA transcripts of (A) myogenin (*MyoG*), (B) serum response factor (*Srf*), (C) striated muscle activator of Rho signaling (*Stars*), (D) α-actin (*Acta1*), (E) myoblast determination protein (*MyoD*), (F) myogenic factor 5 (*Myf5*), (G) creatine kinase, mitochondrial 2 (*Ckmt2*), (H) myosin heavy chain-I (*MyHC-I*), (I) myosin heavy chain-IIa (*MyHC-IIa*), (J) myosin heavy chain-IIb (*MyHC-IIb*), and (K) myosin heavy chain-IIx (*MyHC-IIx*), (L) myosin heavy chain-7 (*Myh7*), (M) tenascin C (*Tnc*), (N) prosaposin (*Pasp*), and (O) frizzled-4 (*Fdz4*) represented as the mean ± SD from three independent experiments and fold change compared between MOCK- and SYNPO2As transduced cells for each day. The numbers below each graph represent the days post-induction (dpi). Statistical significance was determined using two-way ANOVA with post-hoc by Bonferroni. Statistical significance: * p < 0.05, ** p < 0.01, ***p value < 0.005, **** p < 0.001 and NS = Not significant.

## DISCUSSION

First discovered as an actin-binding protein in mouse myoblasts, the short SYNPO2As isoform was originally named myopodin to reflect its preferential expression in skeletal and cardiac muscle, and its similarity to the founding member of the podin family, SYNPO1 (Weins, Schwarz et al. 2001). SYNPO2 interacts with actin and other actin binding proteins, and *in vitro* it nucleates and stimulates actin polymerization and actin bundling (Chalovich and Schroeter 2010, Schroeter, Orlova et al. 2013). In smooth muscle cells, SYNPO2 is a downstream target of SRF and plays a role in regulating expression of smooth muscle differentiation markers (Turczynska, Sward et al. 2015). Our present results indicate the SYNPO2As isoform increases myotube formation during the early stages of myogenesis in skeletal muscle cells and upregulates transcription of a subset of pro-myogenic factors. Taken together, these results suggest a model where the function of SYNPO2As as a regulator of actin polymerization leads to engagement of the SRF and/or STARS actin-regulated signalling pathways to upregulate aspects of the late differentiation pathway and early myotube formation (Fig. 9).

**Figure 9:**
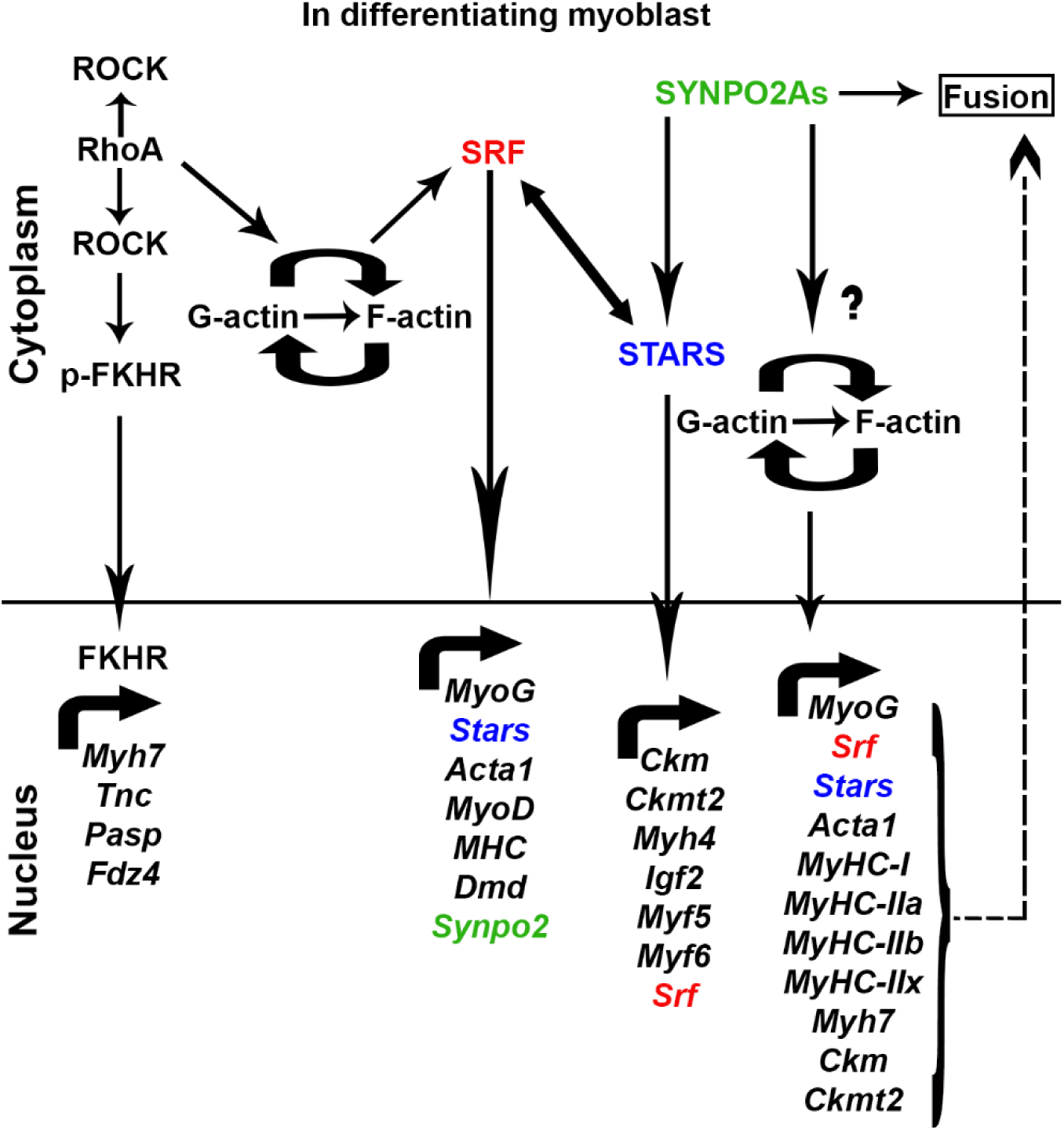
A proposed model of mechanism by which SYNPO2As enhances myoblast fusion. The figure is explained in detail in the text. Briefly, in differentiating myoblasts, the Rho-ROCK pathway is downregulated by RhoE that leads to dephosphorylation of FKHR and nuclear translocation to transcribe myogenic genes, *Myh7, Tnc, Pasp* and *Fdz4*. Down regulated Rho-ROCK pathway also drives the SRF pathway via actin polymerization to transcribe myogenic genes, for example: *MyoG, Stars, Acta1*, *MyoD, MHC, Dmd* and *Synpo2*. Furthermore, STARS, a downstream target of SRF transcribes myogenic genes such as *Ckm, Ckmt2, Myh4, Igf2, Myf5* and *Myf6*. STARS also regulates SRF expression in differentiating myoblasts. SYNPO2As, another downstream target of SRF enhances myogenic genes such as *MyoG, Srf, Stars, Acta1, MyHC-I, MyHC-IIa, b and x, Myh7, Ckm and Ckmt2.* We hypothesize (dotted arrows) that there could be a feedback loop of SRF and STARS to drive the myogenic program, and the enhanced transcript level of the muscle-specific genes could lead to the enhanced fusion phenotype seen in SYNPO2As-transduced C2C12 cells. It remains to be directly demonstrated whether SYNPO2As drives myogenic transcription via actin polymerization.

Expression patterns of the three mouse SYNPO2 isoforms were reflected in a biologically relevant role for SYNPO2As, but not SYNPO2A or SYNPO2B, as a pro-myogenic factor that promotes early myotube formation. The ability to obtain SYNPO2 cDNA from undifferentiated C2C12 myoblast indicated the presence of mRNA transcripts encoding all three mouse SYNPO2 isoforms in pre-differentiated mouse C2C12 myoblast cells. Faint polypeptide bands in western blots of undifferentiated C2C12 cells that co-migrated with SYNPO2As (Fig. 2D) suggested that undifferentiated C2C12 myoblasts may express low levels of SYNPO2. Immunofluorescence microscopy also suggested SYNPO2 is expressed at low levels in newly isolated satellite cells (Fig. 3). Most notably, only expression of the short SYNPO2As isoform was upregulated following differentiation of either cell type, as previously reported in differentiating mouse and human skeletal muscle cells. Expression of the longer SYNPO2 isoforms was not upregulated following differentiation, consistent with the observation that overexpression of these isoforms actually inhibits early myotube formation (Fig. 4). The normal biological function of the long SYNPO2 isoforms appears to be more relevant to muscle tissue maintenance, not muscle development (Ulbricht, Eppler et al. 2013), and the short SYNPO2As isoform (also known as SYNPO2e) is also implicated in myofibril stabilization (Lohanadan, Molt et al. 2021). We now show that the short SYNPO2As isoform promotes the early stages of myogenesis leading to myotube formation.

Some actin-regulatory proteins, such as Rho-GTPases, LIMK1 and mDIA, function as actin polymerizing proteins by binding to G-actin, thereby reducing G-actin levels and activating the SRF pathway (Miralles, Posern et al. 2003). It has been suggested that SYNPO2 may utilize its actin polymerization property to regulate smooth muscle differentiation through this pathway since knockdown of SYNPO2 dramatically reduced the F:G actin ratio and decreased expression of differentiated smooth muscle markers (Turczynska, Sward et al. 2015). The current results provide further evidence in support of this hypothesis. Quantitative RT-PCR analysis revealed that SYNPO2As-transduced cells increased expression of a subset of markers of late myogenic differentiation, primarily those whose expression is regulated by the actin-responsive SRF and STARS pathways. For example, *MyoG*, *Stars* and *Acta1* were enhanced by ∼3-fold and *Srf* by ∼0.2-fold (Figs. 8A-D), while there was no significant increase of *MyoD* levels (Fig. 8E), which implied SYNPO2As was not initiating, but rather selectively enhancing the differentiation program. The transcript levels of *Pasp* and *Fdz4*, two downstream targets in the FKHR pathway, were not affected by SYNPO2As overexpression while two others, *Myh7* and *Tnc*, were significantly enhanced (Fig. 8K-O). However, both *Myh7* and *Tnc* are also SRF targets (Miano, Long et al. 2007, Chiovaro, Chiquet-Ehrismann et al. 2015), so it is unclear whether the increase in these transcripts is caused by SRF or FOXO1a. The other set of genes that were significantly upregulated were downstream targets of the STARS pathway (Wallace et al., 2016). These genes, *Ckmt2* and the myosin isoforms (*MyHC-I*, *MyHC-IIa*, *IIb* and *IIx*), were significantly upregulated ∼3-4-fold before differentiation (0 dpi) and ∼2-3-fold post-differentiation (Figs. 8G-K). The significant increase of these myogenic markers suggested that SYNPO2As engages the actin-responsive SRF and STARS pathways to regulate expression of pro-myogenic factors involved in early myotube formation.

We propose a new functional role for SYNPO2As whereby early myotube formation is enhanced by upregulating expression of pro-myogenic proteins whose expression is controlled through actin-responsive transcription factors. In normal differentiating myoblasts (Fig. 9), downregulation of the Rho-ROCK pathway allows dephosphorylation of FKHR (a.ka. FOXO1a) and nuclear translocation to initiate expression of muscle-specific transcripts such *Myh7*, *Tnc*, *Pasp* and *Fdz4* (Castellani, Salvati et al. 2006, Fortier, Comunale et al. 2008). Another pathway that initiates late myogenic genes is the RhoA-ROCK-SRF pathway. The Rho-ROCK pathway activates SRF via actin polymerization to drive myogenic differentiation (Kuwahara, Barrientos et al. 2005). In differentiating myoblasts, downregulation of the RhoA-ROCK-SRF pathway downregulates MyoD but enhances late differentiation genes such as MHC and α-actin (Iwasaki, Hayashi et al. 2008). In addition to *Synpo2*, the SRF pathway also upregulates many muscle-specific transcripts such as *Acta1, Stars, MyoG, Cnn2, Flna* and *Srf* itself by a feedback loop mechanism (Gauthier-Rouviere, Vandromme et al. 1996, Miano, Long et al. 2007, Randrianarison-Huetz, Papaefthymiou et al. 2018). Another interesting actin adaptor is STARS, which is an actin-binding protein that drives the SRF pathway by polymerizing G-actin (Kuwahara, Barrientos et al. 2005). Overexpression of STARS in C2C12 myoblasts also activates transcription of muscle-specific markers and enhances myoblast fusion (Wallace, Della Gatta et al. 2016). We have shown that overexpression of SYNPO2, an actin polymerizing protein, significantly increased STARS transcript levels∼2-3-fold before and after differentiation (Fig. 8C), and knocking down SYNPO2As reduced STARS transcripts levels by about 30-40% on 1dpi and about 15% on 2dpi (Supplementary Fig. 2G). In addition, overexpressing SYNPO2As also upregulated other muscle specific genes such as *Acta1*, *Stars*, *MyoG*, *Srf*, *Ckm*, *Ckmt2*, *MyHC-1*, *MyHC-IIa*, *MyHC-IIb*, and *MyHC-IIx* (Fig. 9). It remains to be directly determined whether SYNPO2As enhances these muscle-specific transcripts using its actin polymerizing activity to setup a feedback loop in the SRF-STARS pathways. The link between the actin cytoskeleton and gene transcription is a known function of Z-disc associated proteins like SYNPO2.

## EXPERIMENTAL PROCEDURES

### Cells and antibodies

C2C12 (CRL-1772) mouse myoblast cells were obtained from the American Type Culture Collection (ATCC). Phoenix cells were kindly provided by Craig McCormick (Dalhousie University). Cells were grown in Dulbecco Modified Eagle Medium (DMEM) supplemented with 10% fetal bovine serum (FBS) at 37°C with 5% CO_2_. For differentiation experiments, growth medium was replaced with Dulbecco Modified Eagle Medium (DMEM) supplemented with 2% horse serum (HS). Primary antibodies against SYNPO2 (Abcam Ab103710 and Abcam Ab50192), c-myc (Sigma M4439), myosin heavy chain (Developmental Studies Hybridoma Bank MF-20), MyoD (Santa Cruz Biotechnology M-318; Dako M3512), myogenin (Sigma-Aldrich F5D), Pax7 (Developmental Studies Hybridoma Bank), Troponin T (Sigma-Aldrich T6277), LC3-II (3868, Cell signalling), and actin (A2066, Sigma), and secondary antibodies goat-anti-rabbit HRP (474-1506, KPL), goat-anti-mouse HRP (sc-2005, Santa Cruz), and goat-anti-mouse Alexa 488 (Molecular Probes) were purchased from the indicated suppliers. DRAQ5 (Cell Signalling) was used for nuclear staining. 0.1% naphthol blue solution was prepared in-house and used for total protein loading.

### Molecular cloning

RNA was extracted from proliferating C2C12 cells using the RNeasy Mini Kit (Qiagen) as per manufacturer’s instructions and used as a template for cDNA synthesis using an oligo(dT) primer, and cDNA was amplified using isoform specific primers and PfuUltra High Fidelity DNA polymerase (Agilent Biotechnologies). The primers used for PCR amplification flanked the translation start and stop sites in the mouse mRNA sequences for SYNPO2A (Accession #Q91YE8.2), SYNPO2B (Accession #NM_080451.2) and SYNPO2As (Accession #AJ306625.1). Amplicons were cloned into the BamHI-NotI sites in the pBMN retrovirus plasmid vector. All cDNA clones were sequenced in their entirety in both directions, and sequences agreed with those deposited in the databases.

### Generation of stably transduced C2C12 cells expressing SYNPO2 isoforms

Phoenix cells were seeded at 3×10^6^ cells per 100mm dish in DMEM medium containing 25mM HEPES and 10% FBS. Cells were transfected with the empty pBMN retrovirus plasmid vector (kindly provided by Dr. Craig McCormick, Dalhousie University) or with the same vector containing the SYNPO2 cDNAs using poly(ethylenimine) (PEI) transfection reagent. After 48 h, cell culture supernatants were collected and filtered through a 0.45 µM filter to remove cell debris. Sequabrene (4 µg/ml, Sigma) was added to the supernatant to increase the efficiency of viral infection, and C2C12 cells were infected with the retrovirus-containing supernatant for 24 h and then cultured in fresh growth media containing 1µg/ml puromycin (Invitrogen) for 3 days to select pBMN-SYNPO2 transduced cells.

### Generation of knockdown cell lines

Two short hairpin RNAs, one targeting the unique 5’-exon of SYNPO2As (shRNA1) and the other targeting the conserved exon present in all SYNPO2 isoforms (shRNA2), were designed using RNAi Central software (http://cancan.cshl.edu/RNAi_central/RNAi.cgi?type=shRNA) and cloned into the pSMN retroviral plasmid (kindly provided by Dr. Craig McCormick, Dalhousie University). The shRNA2 plasmid construct was transiently transfected into HEK293T cells along with the pBMN plasmids expressing the SYNPO2A or SYNPO2B isoforms, while retroviruses expressing shRNA1 to knockdown endogenous SYNPO2As were used to generate stably transduced C2C12 cells, selected using 2mg/ml G418 (Invitrogen) for 5 days. Western blotting with anti-SYNPO2 antibody was used to determine the knockdown efficiency of the shRNAs.

### Western blotting

Cells were trypsinized, harvested by centrifugation at 500xg for 5 minutes, washed with phosphate buffered saline (PBS), and cell pellets were stored at 80^◦^C until analyzed. Cell pellets were lysed in cell lysis buffer (50 mM Tris, 10 mM MgCl2, 0.5 M NaCl, 2% Igepal, pH 7.5) containing protease inhibitor cocktail (Pierce), sonicated using a stainless-steel probe sonicator, and insoluble debris removed by centrifugation at 15000xg for 10 min. An aliquot of each sample was used to determine protein concentration using the Bio-Rad DC^TM^ protein assay kit, as per manufacturer’s instructions, and the remaining sample was frozen using liquid nitrogen until further processing. Frozen samples were thawed, diluted with cell lysis buffer and protein sample buffer (5% sodium dodecyl sulfate, 0.25% bromophenol blue and 50% glycerol) to equalise the protein concentration, boiled at 95^◦^C for 5 min, and equal protein loads were fractionated by SDS-PAGE (7.5% polyacrylamide). Fractionated samples were transferred onto polyvinylidene difluoride (PVDF) membranes, blocked for 1 h at room temperature using 5% skim milk in TBST (50 mM Tris-HCl, 150 mM NaCl, 0.1% Tween 20), and probed with the respective primary antibodies diluted in TBST at the following dilutions: SYNPO2, 1:2500; MHC, 1:250; myoD, 1:250; myogenin, 1:100; LC3-II, 1:1000; actin, 1:5000. Blots were washed extensively with TBST, treated with horseradish peroxidase-conjugated secondary antibody (1:5000), and developed using ECL-Plus reagent (GE Healthcare) and visualized on a Typhoon 9410 multi-mode imager, or developed using Clarity western ECL substrate (Bio-Rad) and visualized on a Bio-Rad ChemiDoc imaging system. Western blots to examine any changes in the differentiation program were performed in triplicate and relative expression levels were normalized to protein loads detected by staining blots with 0.1% naphthol blue for 30 mins prior to blocking and imaged using the Bio-Rad ChemiDoc calorimetry setting.

### Satellite cell isolation and immunofluorescence microscopy

Care and handling of animals were in accordance with the federal Health of Animals Act, as practiced by McGill University and the Lady Davis Institute for Medical Research. Satellite cells were prepared from abdominal muscles and diaphragms of 8–12-week old *Pax3^GFP/+^* mice by enzymatic dissociation as previously described (Montarras, Morgan et al. 2005). For live cell sorting, single cells were stained with 1 µg/ml propidium iodide (PI) to exclude PI+ dead cells. Cell sorting was performed with a FACSAria Fusion Cell Sorter (BD). Isolated *Pax3^GFP/+^* satellite cells were resuspended in 39% DMEM, 39% F12 (Gibco) growth medium containing 20% FBS (Gibco) and 2% Ultroser G (Pall, Port Washington, NY). Cells were cultured in 35-mm dishes coated with 0.2% gelatin at a density of 7500 cells per dish. For immunocytochemical analysis, cultured cells were fixed with 10% formalin and permeabilized with 0.2% Triton X100, 50 mM NH_4_Cl in PBS. Cells were incubated with 0.2% fishskin gelatin in PBS (Sigma). The following antibodies were used as primary antibodies: anti-MyoD (1:100, Santa Cruz Biotechnology, Santa Cruz, CA), anti-Troponin T (1:300, Sigma) and anti-SYNPO2 (1:100, Abcam Ab103710). Secondary antibodies were coupled to flurochromes Alexa 488 or 594 (1∶500, Molecular Probes, Carlsbad, CA). 4,6-Diamidino-2-phenylindole (DAPI, 1:500, Molecular Probes) was used to counter-stain nuclei. All images were captured at the same microscope settings so that fluorescence intensity was representative of detection levels.

### Indirect immunofluorescence microscopy

To observe endogenous SYNPO2 subcellular localization, C2C12 cells at 80% confluence grown on coverslips were induced to differentiate using medium containing 2% horse serum until 2 dpi. Cells were fixed with 4% paraformaldehyde, permeabilized with 0.25% triton X-100 and blocked with 1% BSA at room temperature. Cells were then incubated with anti-SYNPO2 antibody (1:500) overnight at 4◦ C. Cells were washed and incubated with goat anti-rabbit secondary antibody (Alexa fluor 488), and Alexa-555 conjugated phalloidin for 1 h at room temperature, and cell nuclei were stained using DRAQ5 (1:1000) for 10 minutes. A similar protocol was used for detecting ectopically expressed myc-tagged SYNPO2 isoforms using anti-c-myc primary antibody. A Z-stack of the cells were imaged using a 63X 1.4 NA oil-immersion objective lens on a using a Zeiss LSM 510 Meta laser scanning confocal microscope. Images were acquired using Zen software (Zeiss) and a single slice from each Z-stack was used. The orthogonal view of the Z-stack was obtained by selecting the slice of interest and the images were processed using Photoshop CS5 (Adobe).

### Quantification of myotube formation

Duplicate or triplicate wells in a 12-well plate were seeded with C2C12 cells at 80% confluence, cultured in growth medium for 24 h, and then induced to differentiate using medium containing 2% horse serum until 3-4 dpi. Cells were fixed with methanol for 15 minutes and stained with Wright-Giemsa stain (Siemens). A total of 10-12 random fields from each well were imaged using a 10x objective. For the actin inhibitor studies, 10µM ROCK-inhibitor (Y-27632) or 20µM Arp2/3 inhibitor (CK666) were added to cells 8 h after differentiation (to ensure the inhibitors did not affect the differentiation program), and media containing the inhibitors or solvent control was replaced every 24 h until 3dpi. Cells were methanol fixed, immunostained with anti-MHC and Hoescht, and imaged at 10X using the EVOS FL cell imaging system (Thermofisher). All experiments were repeated three times and the fusion index was quantified as the percent of the total number of nuclei in a field present in MHC+ myotubes containing 3 or more nuclei to the total number of nuclei present in a field. All results are presented as the mean ± SD of the three experiments.

### Quantitative real-time PCR

C2C12 myoblasts stably transduced with mock or SYNPO2As, or control ShRNA or SYNPO2 ShRNA (Sh2) constructs were seeded in a 12-well plate in triplicate and RNA was extracted at 0 to 3 dpi. RNA was extracted using the RNeasy mini kit (#74106,Qiagen) according to the protocol. Briefly, 350 μl of RLT buffer was added to each well and cell lysate was pooled from the three wells and homogenized using a 20- gauge needle. One volume of 70% ethanol was added to the homogenized sample, mixed well and centrifuged in a gDNA eliminator spin column to remove gDNA contamination. The elute was washed with 700ul of RW1 buffer followed by RPE buffer and eluted in RNase and DNase free water. cDNA synthesis was carried out using RNA to cDNA EcoDryTM Premix kit (Takara Bio # 639549). Briefly, 2 μg of RNA was added to the mix and incubated at 42◦C for 1 h, and reaction was stopped by incubating at 70◦ C for 10 minutes. A standard curve was performed for primer validation. The synthesized cDNA was diluted 1:2 and used for qRT-PCR analysis. The qRT-PCR was performed using FastStart Essential DNA green master (Roche, # 06402712001). The primers used in this study are detailed in Table 1.

**Table 1.**
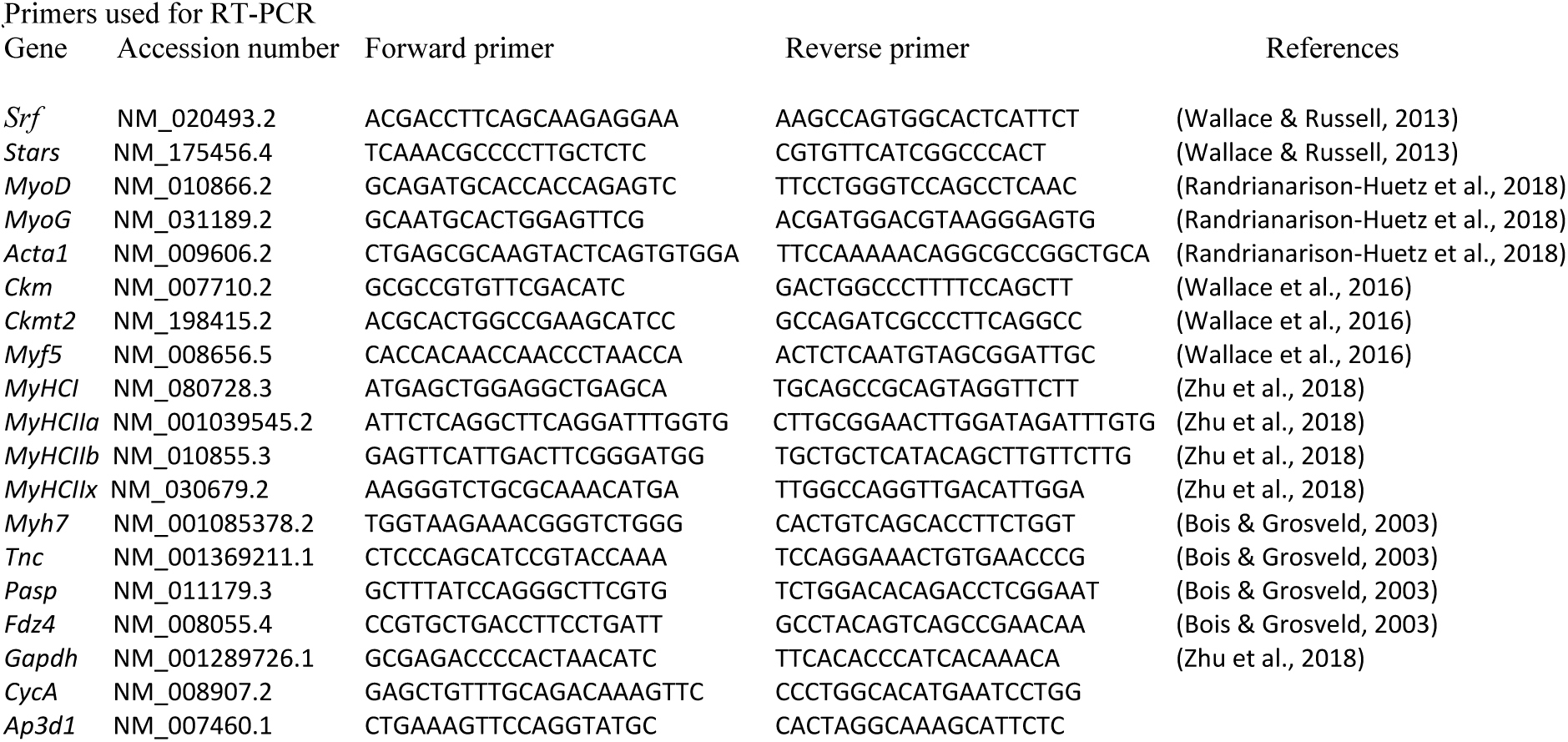
Primers used for RT-PCR.

### Statistical analysis

All results reported herein are the mean ± SD of duplicate or triplicate experiments as indicated. Statistical significance was determined using an unpaired Student *t* test (p < 0.05), or one-way or two-way ANOVA along with post-hoc by Bonferroni with a 95% confidence interval as indicated in the figure legends.

## ACKNOWLEDGMENTS

R.D. was supported by grants from the Canadian Institutes of Health Research (CIHR) and the Natural Sciences and Engineering Research Council of Canada (NSERC). N.M. was supported by a Nova Scotia Graduate Scholarship. F.K. was supported by a scholarship from the Cancer Research Training Program (CRTP) with funding from the Dalhousie Cancer Research Program (DCRP) and the Dalhousie Medical Research Foundation (DMRF). C.C. was supported by grants from the CIHR, the Muscular Dystrophy Association, and the Stem Cell Network. Thanks to all the staff of the Cellular and Molecular Digital Imaging facility at Dalhousie University.

