## Supplementary Figures for "The Short Isoform of the Mouse Actin Adaptor Protein Synaptopodin-2 Activates Actin-Responsive Transcription Factors and Enhances Myoblast Fusion"

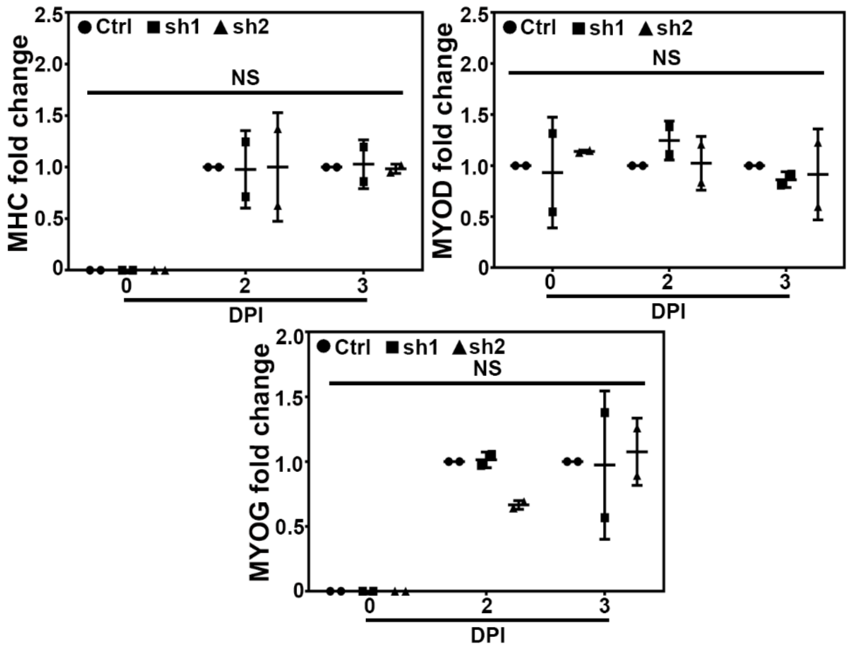


**Supplementary Figure 1: SYNPO2 isoform knockdown does not alter the expression level of myogenic differentiation proteins.** Quantified data of MHC, MYOD and MYOG western blots represented as the mean ± SD from two independent experiments of control, shRNA1 (sh1) and shRNA2 (sh2) cell lysates collected at 0, 2, 3 dpi and probed with anti-MHC, MYOD or MYOG antibody. Statistical significance was determined using two-way ANOVA with post-hoc by Bonferroni. Statistical significance: NS = Not significant.

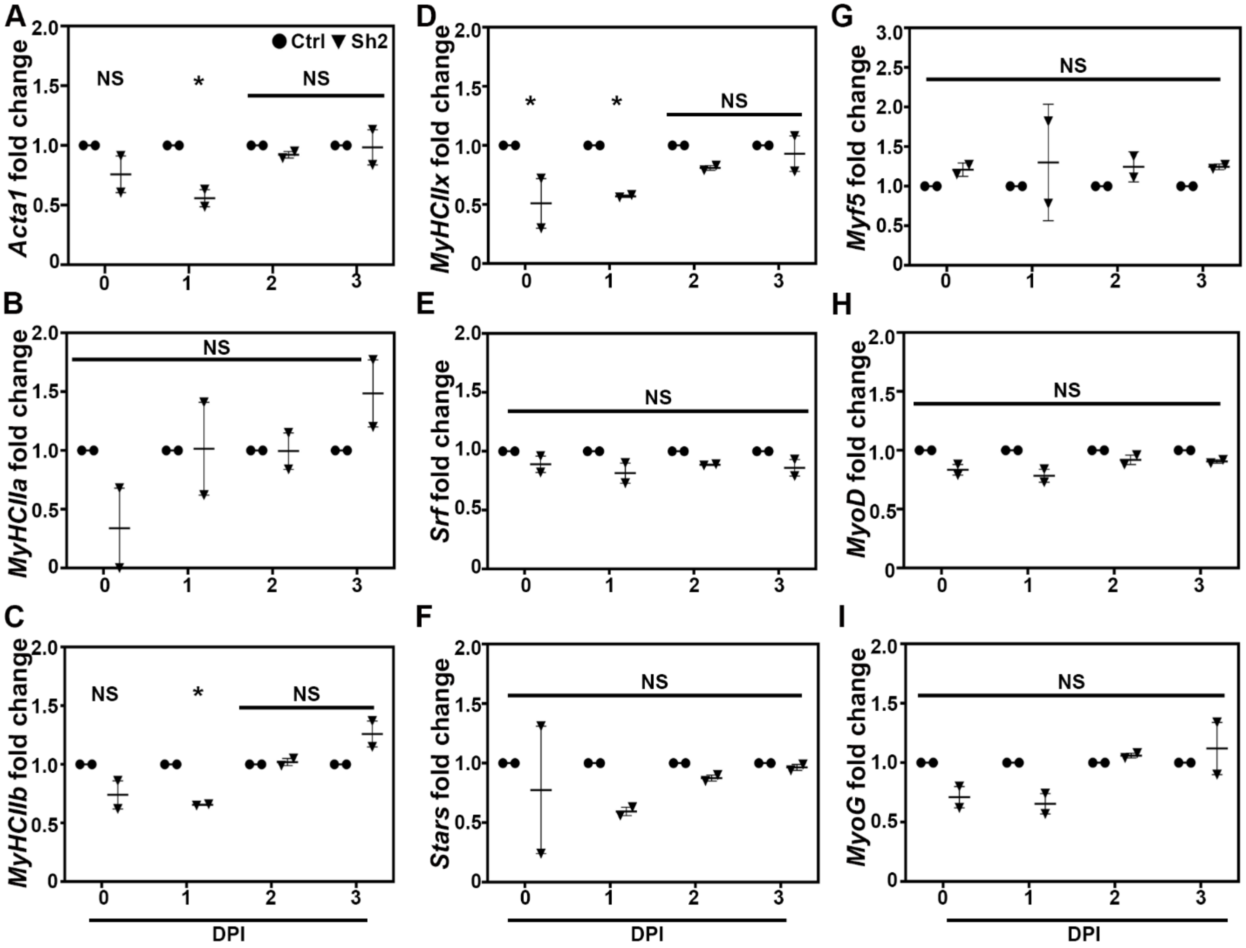


**Supplementary Figure 2: SYNPO2 isoforms knockdown decreased transcript level of selective myogenic genes.**
RT-qPCR of selective myogenic genes mRNA transcripts of (A) α-actin (*Acta1*), (B) myosin heavy chain-I (*MyHC-I*), (C) myosin heavy chain-IIa (*MyHC-IIa*), (D) myosin heavy chain-IIb (*MyHC-IIb*), (E) myosin heavy chain-IIx (*MyHC-IIx*), (F) serum response factor (*Srf*), (G) striated muscle activator of Rho signaling (*Stars*), (H) myogenic factor 5 (*Myf5*), (I) myoblast determination protein (*MyoD*), (J) myogenin (*MyoG*), (K) frizzled-4 (*Fdz4*), and (L) creatine kinase, mitochondrial 2 (*Ckmt2*) are represented as the mean ± SD from two independent experiments and fold change compared between Ctrl and knockdown (Sh2) cells for each day. The numbers below each graph represent the days post-induction (dpi). Statistical significance was determined using two-way ANOVA with post-hoc by Bonferroni. Statistical significance: * $p < 0.05$ and NS = Not significant
